# Reconciling GenBank names with standardized avian taxonomies to improve linkage between phylogeny and phenotype

**DOI:** 10.1101/2022.02.07.479408

**Authors:** Peter A. Hosner, Min Zhao, Rebecca T. Kimball, Edward L. Braun, J. Gordon Burleigh

## Abstract

Biodiversity research has advanced by testing expectations of ecological and evolutionary hypotheses through the linking of large-scale genetic, distributional, and trait datasets. The rise of molecular systematics over the past 30 years has resulted in a wealth of DNA sequence data from around the globe, facilitating biodiversity research. However, advances in molecular systematics also have created taxonomic instability, as new estimates of evolutionary relationships and interpretations of species limits have led to widespread scientific name changes. Taxonomic instability, or “splits, lumps, and shuffles”, present logistical challenges to large-scale biodiversity research because species or populations may be listed under different names in different data sources, or because different species or populations may be listed under previous names. Consequently, distributional and trait data are often difficult to link directly to DNA sequence data without extensive and time consuming curation. Here, we present RANT: Reconciliation of Avian NCBI Taxonomy. RANT applies taxonomic reconciliation to standardize all avian names in use in NCBI GenBank, a primary source of genetic data, to a widely-used and regularly-updated avian taxonomy: eBird/Clements. Of 14,341 avian species or subspecies names used by GenBank, 11,031 names directly matched an eBird/Clements name, which were linked to over 6 million nucleotide sequences. For the remaining unique avian names in GenBank, we used Avibase’s taxonomic concepts, taxonomic descriptions in Cornell’s Birds of the World, and DNA sequence metadata to identify corresponding eBird/Clements names. Reconciled names were linked to over 600,000 nucleotide sequences, approximately 9% of all avian sequences on GenBank. Nearly 10% of eBird/Clements names had nucleotide sequences listed under two or more GenBank names. Our avian GenBank naming reconciliation is open source and available at GitHub, where it can be updated to correspond with future annual eBird/Clements taxonomic updates.

**LAY SUMMARY:**

– 23% of avian names on GenBank do not match eBird/Clements, a widely-used standardized avian taxonomy
– 600,000 nucleotide sequences on GenBank are associated with names that do not match eBird/Clements
– 10% of eBird/Clements names have nucleotide sequences listed under multiple GenBank names
– We provide an open source taxonomic reconciliation to mitigate difficulties associated with non-standardized name use for GenBank data

## INTRODUCTION

Public data repositories are rich information sources constituting vital infrastructure for integrative and large-scale research in organismal biology. As a taxonomic group, birds are well-suited to these endeavors. Their global ubiquity, relative ease of observance and identification, and charismatic appearances lend to their enduring popularity among professional and recreational scientists alike. The quantity and extent of avian data has proliferated in recent years, a direct result of efforts to grow and share these data (Table 1). The information available documenting and describing avian genetics, population dynamics, distributions, behaviors, and physical traits has become truly staggering.

**Table 1.**
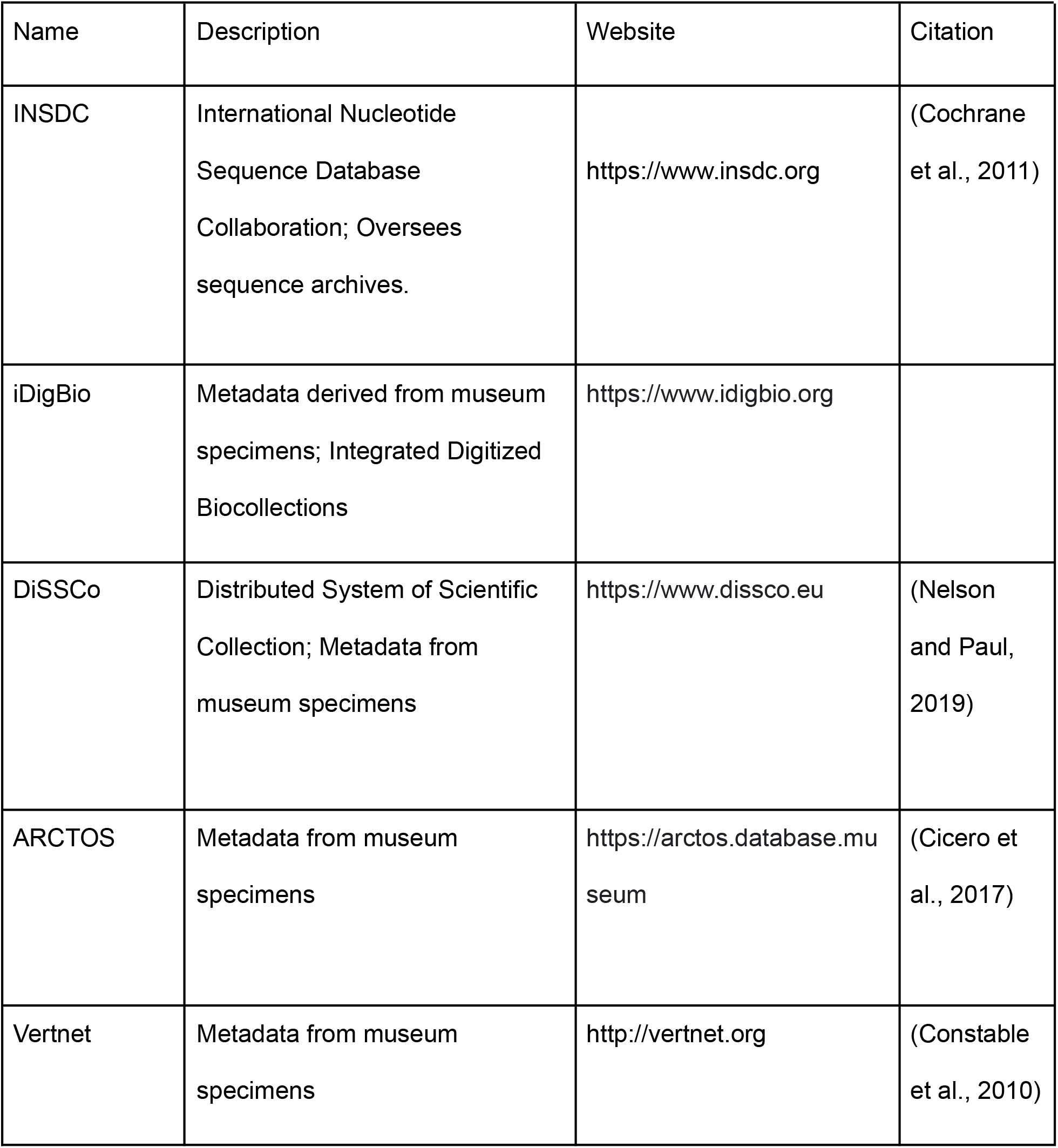
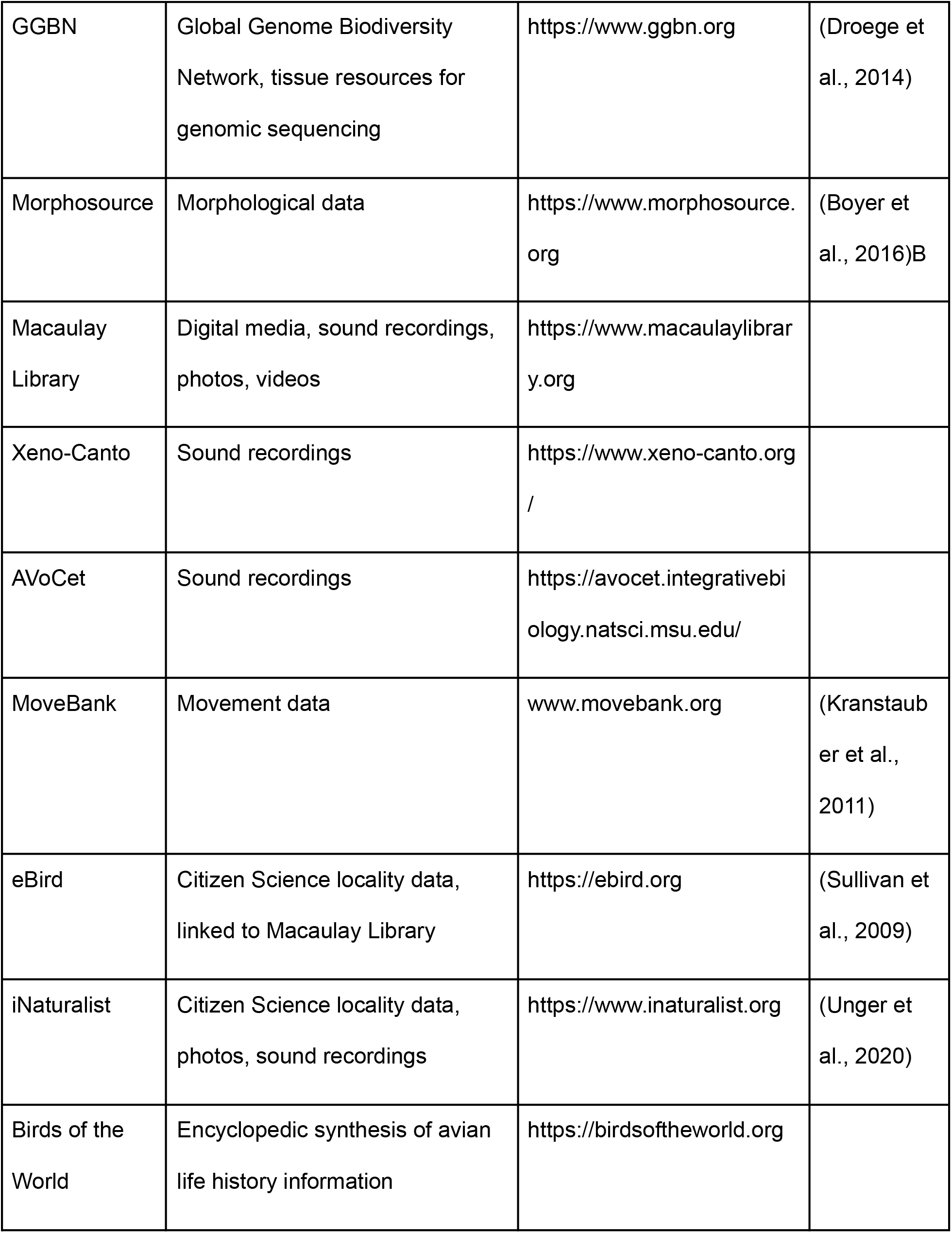
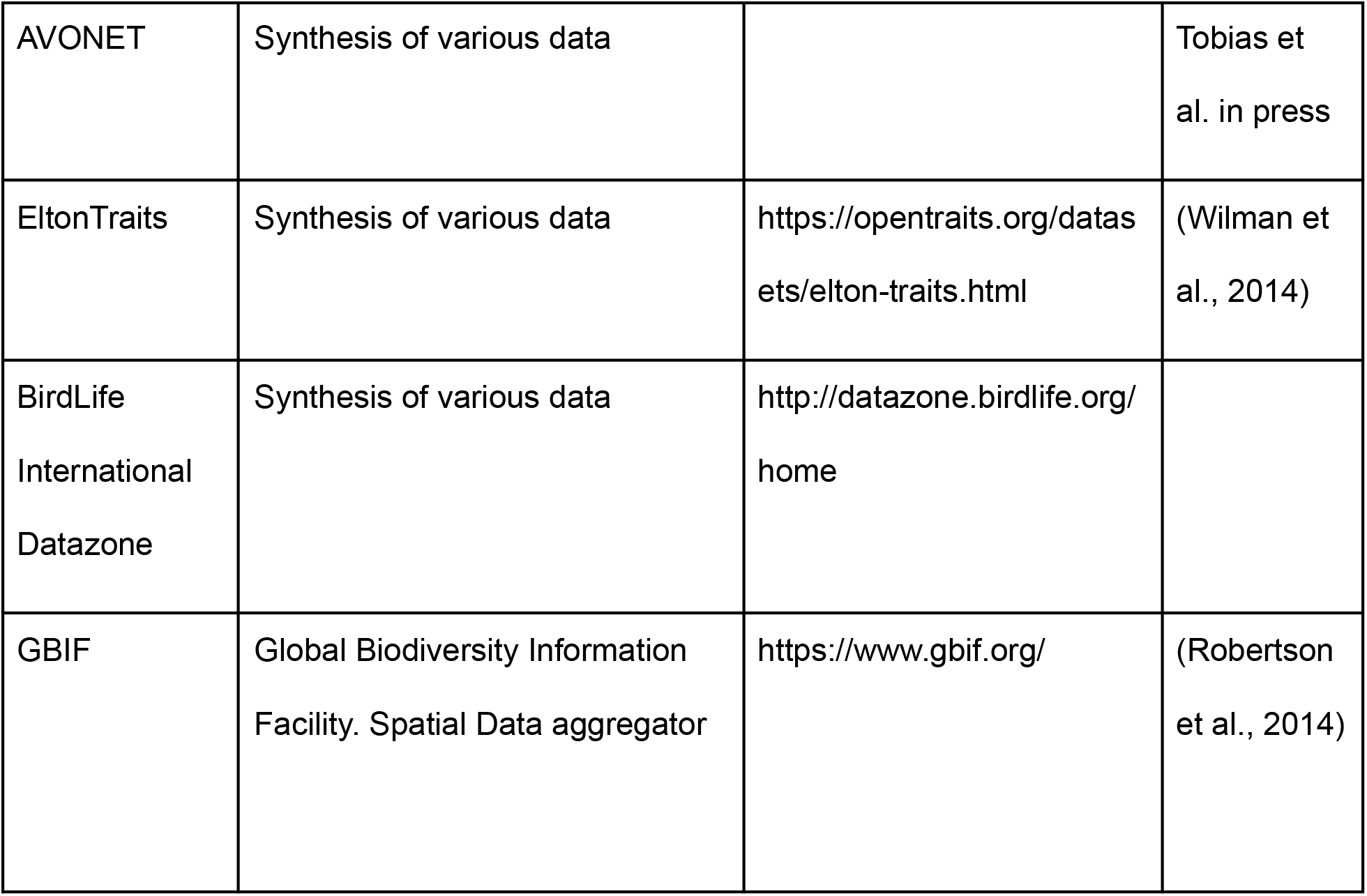
Examples, descriptions, links, and references to large global databases that contain avian data.

| Name | Description | Website | Citation |
| --- | --- | --- | --- |
| INSDC | International Nucleotide Sequence Database Collaboration; Oversees sequence archives. | https://www.insdc.org | (Cochrane et al., 2011) |
| iDigBio | Metadata derived from museum specimens; Integrated Digitized Biocollections | https://www.idigbio.org |  |
| DiSSCo | Distributed System of Scientific Collection; Metadata from museum specimens | https://www.dissco.eu | (Nelson and Paul, 2019) |
| ARCTOS | Metadata from museum specimens | https://arctos.database.museum | (Cicero et al., 2017) |
| Vertnet | Metadata from museum specimens | http://vertnet.org | (Constable et al., 2010) |
| GGBN | Global Genome Biodiversity Network, tissue resources for genomic sequencing | https://www.ggbn.org | (Droege et al., 2014) |
| Morphosource | Morphological data | https://www.morphosource.org | (Boyer et al., 2016)B |
| Macaulay Library | Digital media, sound recordings, photos, videos | https://www.macaulaylibrary.org |  |
| Xeno-Canto | Sound recordings | https://www.xeno-canto.org/ |  |
| AVoCet | Sound recordings | https://avocet.integrativebiology.natsci.msu.edu/ |  |
| MoveBank | Movement data | www.movebank.org | (Kranstauber et al., 2011) |
| eBird | Citizen Science locality data, linked to Macaulay Library | https://ebird.org | (Sullivan et al., 2009) |
| iNaturalist | Citizen Science locality data, photos, sound recordings | https://www.inaturalist.org | (Unger et al., 2020) |
| Birds of the World | Encyclopedic synthesis of avian life history information | https://birdsoftheworld.org |  |
| AVONET | Synthesis of various data |  | Tobias et al. in press |
| EltonTraits | Synthesis of various data | https://opentraits.org/datasets/elton-traits.html | (Wilman et al., 2014) |
| BirdLife International Datazone | Synthesis of various data | http://datazone.birdlife.org/home |  |
| GBIF | Global Biodiversity Information Facility. Spatial Data aggregator | https://www.gbif.org/ | (Robertson et al., 2014) |

To leverage the vast wealth of avian information and effectively implement phylogenetic comparative methods (Felsenstein, 1985) and other evolutionary analyses, it is crucial to have clear, one-to-one linkage between data records and the populations of organisms to which they are derived. Over the past 30 years, molecular systematics has wholly transformed avian taxonomy and nomenclature (Barrowclough et al., 2016; Gill, 2014; Sangster, 2009). Its insights have reorganized the avian tree of life (Beresford et al., 2005; Braun et al., 2019; Hackett et al., 2008; Harvey et al., 2020; Jarvis et al., 2014; Lovette et al., 2010; Moyle et al., 2012; Oliveros et al., 2019) and reformed practical applications of species limits (Andersen et al., 2014; Hosner et al., 2018). An unfortunate consequence of these much-needed reorganizations is that they often require changes to organisms' scientific names. In modern implementations of the Linnaean system of nomenclature, higher taxa (e.g., genus, family, order) are required to be monophyletic. Hence, any move of a species to a different genus, or an update of species limits, requires scientific name changes for populations.

Identifying and tracking avian nomenclatural changes over time is itself a difficult task. As an example, we compared two major taxonomic works completed before DNA sequencing technology was widely available in ornithology, the Peters checklist series (1931–1987; (Bock and Paynter, 1990) and Sibley and Monroe (1993), to the the eBird/Clements (2019) list (Data Repository D1). Only 6288 of the 9204 (69%) Peters checklist species names, and 7470 of 9702 (77%) of the Sibley and Monroe (1993) species names matched exactly to the 10721 eBird/Clements (2019) names. Allowing the last two letters of the species epithet to mismatch, to account for minor differences in spelling, only improved name matching slightly (27 more matches for the Peters checklist, 20 more matches for Sibley & Monroe 1993). Although the details of broad list comparisons will vary depending on exactly which taxonomies are compared, all modern avian taxonomies differ substantially from corresponding works produced only decades ago.

In addition to instability stemming from name changes through time, another contributor to scientific name instability is the presence of multiple competing standardized avian taxonomies. Currently, there are four main global choices: eBird/Clements (Clements et al., 2019), IOC World Bird List (Gill et al., 2021), Howard & Moore Complete Checklist of Birds of the World (Dickinson and Christidis, 2013; Dickinson and Remsen Jr., 2014), and HBW/BirdLife Taxonomic Checklist (Burfield et al., 2017). Although similar in many respects, each of these lists are governed differently, are updated at different intervals, and apply species recognition criteria differently (Garnett and Christidis, 2017). For example, in raptors, a paraphyletic assemblage of predatory non-passerine landbirds, McClure et al., (2020) found that major world lists disagreed in species-level name application in 11–25% of cases. Beyond these most-referenced world lists, there are additional regional and country specific avian taxonomies.

Different biodiversity databases often use different underlying taxonomies, requiring users to reconcile names between sources (Boyle et al., 2013; Lepage et al., 2014) before downstream analyses are prudent. Some large avian data sources employ standardized global avian taxonomies from the start. For example, eBird (https://www.eBird.org/) and the Macaulay Library (https://www.macaulaylibrary.org/) use the related eBird/Clements taxonomy, which is usually updated annually. On the other hand, Xeno-canto (https://www.xeno-canto.org/), uses the IOC World Bird List, which is updated up twice a year— although it has been updated quadrennially in the past. Using standardized taxonomies for databases vastly improves the ability for users to identify discrepancies between name usage and application, especially through “taxonomic concepts” as implemented in Avibase (Lepage et al., 2014; McClure et al., 2020). Name reconciliation can be trivial when working only with a few familiar taxa, but it requires extraordinary time and effort when managing large numbers of taxa and when working at global scales. Taxonomic reconciliation becomes even more difficult and time-consuming when data sources implement their own taxonomy *de novo* in lieu of a standardized list, or when datasources lack consistent name use. For databases where the taxonomic names are not readily traceable, it can be impossible to correctly link information from one database to another without supplemental information. Failure to correctly link names may cause available information to be ignored, excluded, or worse— that data are mis-attributed to the wrong population (McClure et al., 2020). This issue could be particularly problematic for bird groups in geographic regions disproportionately affected by taxonomic progress (Neate-Clegg et al., 2021), or for poorly-known birds with limited data such as rare or endangered taxa. In some cases, opportunities to better understand these regions and their birdlife could be lost simply because of taxonomic instability.

NCBI GenBank (Benson et al., 2012), a partner of the INSDC, is the major data repository and distributor for biodiversity genetic data used in phylogenetic analyses. Accurate phylogenetic inference underpins most modern comparative studies, and hence it is necessary to confront naming issues in GenBank data before assembling large-scale, synthetic phylogenies (Burleigh et al., 2015; Jetz et al., 2012) and before linking such phylogenies to other comparative datasets (Pigot et al., 2018). Although GenBank implements policies to standardize names (Schoch et al., 2020), it does not rely on any single standardized avian taxonomy. GenBank policy states that taxonomic names must be published and valid, but in practice names are user-submitted and sometimes informal. Furthermore, as names are updated and changed by some or all standardized avian taxonomies, GenBank largely relies on the original data uploader to curate and update records. This can lead to problems in light of taxonomic instability. For example, when the name of a species changes (e.g., moved to a new genus, or a different specific epithet is used), sequences may be organized under both former and present names. Hence, a researcher may obtain some sequences for a given taxon, but may not realize that other sequence data exists. Worse, a user may assume no data exists for a given taxon, as it could be listed under a former name without current acceptance in standardized lists. Additional uncertainties arise when species are split into two or more entities, or when species are lumped yet remain listed in GenBank under multiple names.

Ultimately, the only way to link GenBank sequences with other types of comparative data is to reconcile GenBank’s avian names to standardized avian taxonomies. One strategy is producing an open source, parallel data structure, which can be curated and updated as avian taxonomy changes (Leray et al., 2020; Riginos et al., 2020). Each GenBank name has a unique numerical identifier (TaxID; Schoch et al., 2020) and each GenBank database record has a unique identifier. Using these identifiers, it is possible to link one or many names, corresponding to standardized lists or to Avibase taxonomic concepts. Here, we attempt such a reconciliation, linking GenBank taxon identifiers to the eBird/Clements 2019 list for all avian GenBank TaxIDs. To further explore the extent to which taxonomic instability and its biases affect birds, we summarize avian data patterns related to taxonomic groups, geographical areas, and conservation status. Finally, we summarize the extent to which name-reconciled sequences apply to large comparative databases, namely Macaulay Library and Xeno-canto bird sound vocalizations, using the GenBank Nucleotide database, the GenBank product with the broadest taxonomic coverage.

Our goal was to reconcile the taxonomic names in GenBank (TaxIDs) to a major avian taxonomy in order to link GenBank sequences, and phylogenetic trees built from these sequences, to ancillary data sources. We selected eBird/Clements v2019 as the focal standardized bird list, because of its use in the world's largest bird observation dataset (eBird), its related media resources (Macaulay Library), and its linked Birds of the World information content. Existing tools can reconcile the Clements list with other standardized taxonomies (Gill et al., 2021; Lepage et al., 2014). Hence, once GenBank names are linked to a single standardized taxonomy, in this case eBird/Clements, reconciling to other standardized taxonomies (IOC, BirdLife International, Howard & Moore) is straightforward.

## METHODS

### Taxonomic reconciliation

We downloaded all names from the NCBI Taxonomy database (Schoch et al., 2020) that descended from “Aves” (TaxID: 8782) on 3 May 2020 (Data Repository D2). From this list, we extracted all species and subspecies names as well as their NCBI Taxonomy ID (TaxID) numbers. We then ran a custom Perl script (Data Repository D3) to exactly match binomial (genus, species) and trinomial (genus, species, subspecies) names from NCBI Taxonomy to the names recognized by eBird/Clements v2019 Integrated Checklist (August 2019; Data Repository D4). For each mismatch with the NCBI Taxonomy name, we then identified the corresponding equivalent eBird/Clements species or subspecies. We first searched for names in Avibase (Lepage et al., 2014). However, Avibase’s search function currently facilitates only exact matches to taxonomies it implements. For names that were not an exact match to an Avibase taxonomic concept, we implemented web searches (Google) which often identified minor spelling differences, consulted Cornell’s Birds of the World Online (https://birdsoftheworld.org), and consulted relevant literature— often the papers that first published those sequence data.

We classified nine categories of naming mismatches resulting from discrepancies between GenBank and eBird/Clements names (Table 2). We summarized the total number and proportion of reconciled GenBank TaxIDs by bird orders and within the largest bird order Passerformes, by families. We also summarized the number of GenBank nucleotide sequences and number of reconciliations for each IUCN conservation status category. For taxon that did not have a direct match to an IUCN name, we placed it under “Not Assessed”.

**Table 2.** Categories, descriptions, and examples of name mismatches between GenBank and eBird/Clements names.

| Name | Description | Example |
| --- | --- | --- |
| **New** | A species or subspecies that was undescribed when its sequences were uploaded to GenBank. To preserve nomenclature priority, GenBank avoids unpublished or in press names of undescribed taxa, instead assigning an informal placeholder name. Typically, | Megascops_sp._SMD-2015 (TaxID: 1740173) corresponds to the Santa Marta Screech-Owl *Megascops gilesi* Krabbe, 2017. |
|  | the placeholder name consists of the genus, the data uploaders initials, and the year of first upload. |  |
| **Lump** | A name that corresponds to species rank in GenBank, but a subspecies rank in eBird/Clements. | GenBank name *Megascops colombianus* (TaxID: 1740167) corresponds to *Megascops ingens colombianus* in eBird/Clements |
| **Split** | A name that corresponds to a subspecies rank in GenBank, but a species rank in eBird/Clements. | GenBank subspecies name *Otus megalotis everetti* (taxiid: 56274) corresponds to the species name *Otus everetti* in eBird/Clements. |
| **Shuffle** | A taxon that has an equivalent rank in GenBank and eBird/Clements, but different name usage. Most often shuffles stem from changes in genera, but a few species epithets have changed because of new evidence regarding nomenclature priority. | GenBank name *Mimizuku gurneyi* (id: 56287) corresponds to *Otus gurneyi* in eBird/Clements. |
| **Spelling** | A taxon that has an equivalent name in GenBank and eBird/Clements, but a slightly different spelling is implemented. | GenBank name *Glaucidium nanum* (TaxID: 126809) corresponds to the eBird/Clements name *Glaucidium nana*. |
| **Hybrid** | A hybrid individual and usually identified in GenBank by a name comprising the putative parental species separated by a cross "x". | GenBank name *Strix occidentalis* x *Strix varia*.<br>Hybrids were not reconciled to eBird/Clements names, although eBird taxonomy does include and organize names for some frequent avian hybrid parental combinations. |
| **Extinct** | An extinct taxon that is not regulated by eBird/Clements because it was not documented in the modern era. | *Aepyornis maximus* (TaxID: 748142) is known from Holocene bones and eggshell materials that have yielded sequenceable DNA, but this name is not regulated by eBird/Clements |
| **Domesticated** | A domesticated breed or line. | GenBank has a listing for the domesticated “Society Finch” as *Lonchura striata domestica* (TaxID: 299123), but in eBird/Clements it refers to *Lonchura striata* because domesticated forms are not generally considered valid subspecies. |
| **Unidentified** | Refers to TaxIDs where we were unable to assign a species name. | Generally samples not identified to species, or environmental DNA samples. |

### GenBank sequences associated with avian names

We tallied the number of core nucleotide sequences in GenBank associated with each taxonomic ID by downloading the “nucl_gb.accession2TaxID” file on 2 November 2020 (Data Repository D5). This file lists the accession number for each sequence in the GenBank nucleotide database and its corresponding taxonomic ID number. From this, we wrote a Perl script (Data Repository D6) to count the number of nucleotide sequences associated with each taxonomic ID corresponding to an avian taxonomic IDs. To obtain counts of the number of runs in the NCBI Sequence Read Archive (SRA) associated with each bird species, we downloaded the “RunInfo” for the SRA runs (“SraRunInfo.csv”) within “Aves” on August 1, 2021 (Data Repository D7).To obtain counts of the number of genome sequences in GenBank associated with each name, we downloaded from NCBI on September 5, 2021 a summary of the NCBI Genome files (“genome_result.txt”) within “Aves” (Data Repository D8).

### Linking eBird/Clements names to geographic realms

For taxa that were successfully assigned to eBird/Clements species names (either by direct name match or taxonomic reconciliation), we delimited their geographic realms using the associated IOC breeding ranges (eight terrestrial realms and four oceanic realms). Here we implemented IOC, rather than eBird/Clements geographic information because eBird/Clements does not summarize species occurrence by geographic realm. We also manually assigned geographic realms for species without range information available in the IOC v10.1 checklist (master_ioc_list_v10.1.xlsx). We defined species that occur in only one realm as realm endemics, and species that occur in two or more realms as widespread. We then summarized the number of reconciliations and the number of GenBank nucleotide sequences for each realm, and widespread species.

### Linking eBird/Clements names to other databases

Since Macaulay Library uses eBird/Clements taxonomy for its bird images, audios and videos, we can readily link these media resources to the GenBank nucleotide data under the same eBird/Clements names. We downloaded a summary of available media data by April 2021 from Macaulay Library (https://www.macaulaylibrary.org/resources/media-target-species/; Data Repository D9) and used audio data as an example to examine the extent to which name-reconciled GenBank sequences apply to large comparative databases. We also examined a second global avian vocalization database, Xeno-canto, which uses the IOC taxonomy. To match Xeno-canto’s 10,909 avian names to eBird/Clements names, we filtered out the species with a direct name match and then reconciled the remaining using Avibase taxonomic concepts. Lastly, we summed up the number of Xeno-canto sound recordings (by October 2020, https://www.xeno-canto.org/collection/species/all; Data Repository D10) under the same eBird/Clements name. For example, the Xeno-canto name *Colinus leucopogon* had 26 sound recordings and *Colinus cristatus* had 57, but the eBird/Clements name *C. cristatus* would have 83, because *C. leucopogon* is treated as a subspecies of *C. cristatus* by eBird/Clements.

## Results

### Descriptive statistics of taxonomic reconciliation

Of 14,341 GenBank species and subspecies TaxIDs within Aves, we were able to exactly match an eBird/Clements name for 11,031 (77%; Fig. 1; Data Repository D11). Of the 3,310 GenBank names without an exact match, we were able to reconcile 2917 to eBird/Clements names using Avibase taxonomic concepts and other sources. Twenty-three percent of GenBank names needed reconciliation to match with eBird/Clements names, and of non-exact-matching names, we were able to reconcile 88%. By far, the most frequent cause of discrepancy between GenBank and eBird/Clements names were “shuffles” (64%), most often because of a genus name change. Splits (11%) and lumps (11%), owing to classification differences at species/subspecies ranks, were nearly equally frequent. Spelling discrepancies (5%), names of extinct taxa not regulated by eBird/Clements (4%), and hybrids (3%) were relatively infrequent. Finally, only a few new species names (0.7%) or names used for domestic breeds (0.2%) contributed to naming discrepancies. In total, we were unable to assign 393 (3%) of GenBank names to eBird/Clements names.

**Fig 1.**
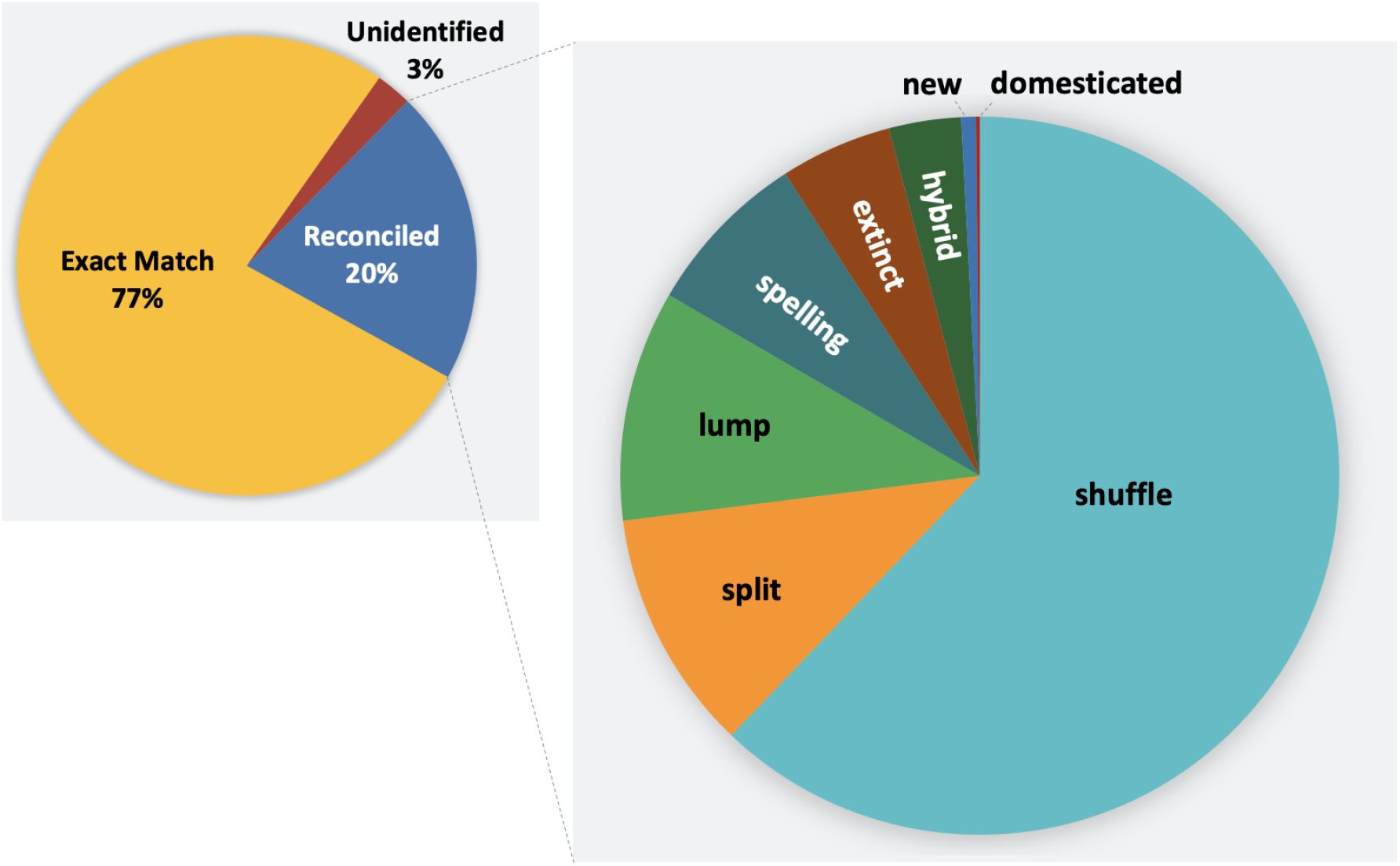
Proportions of GenBank taxa names that are directly matched to eBird/Clements names (Exact Match), manually reconciled to eBird/Clements names (Reconciled), and unidentifiable that includes taxa not identified to the species level or erroneous taxa (Unidentified).

Following reconciliation, we found that 9361 eBird/Clements species names had at least one GenBank Nucleotide sequence attributed, whereas 1832 species had no attributable sequences. We also found that 1050 (10%) of eBird/Clements species names have sequences listed under two or more GenBank names. For the GenBank SRA (sequence read archive, sets of DNA sequence reads derived from massively parallel sequencing runs), 24% of avian species and subspecies were associated with a record. Of the 3375 species and subspecies with SRA data, only 316 (9%) required reconciliation. Among reconciled names, the most common reason was due to shuffles (59% of reconciliations). While many reconciliation categories showed similar proportions to the GenBank data, reconciled names associated with SRA data included a greater proportion of hybrids (7%), domestics (2%), and unidentified (9%), but a lower proportion of splits (3%). Fewer than 4% (20 out of 530) GenBank genome assemblies required reconciliation.

When organized by the number of sequences affected by taxonomic reconciliation, different patterns emerged. In the GenBank Nucleotide database, 6,302,287 (91%) of sequences were a direct match, 626,079 (9%) we reconciled to eBird/Clements, and 2575 (0.02)% we failed to reconcile. Of the nucleotide sequences we reconciled to eBird/Clements, 106,940 (17%) we attributed to shuffles, 16,129 (2.6%) we attributed to lumps, and 381,652 (61%) we attributed to splits. We attributed 1952 sequences to extinct species names not regulated by eBird/Clements, 5909 (0.9%) sequences to hybrids, 102 (0.016%) sequences to new species names, and 110,748 (17%) sequences to domestic breeds.

The total number and proportion of sequences reconciled varied substantially among bird orders and among families within Passeriformes (Figs. 2 & 3). Orders with the largest numbers of reconciled taxa corresponded to those with the greatest species diversity, including the Passeriformes (Songbirds), Piciformes (Woodpeckers and allies) and Caprimulgiformes (nightjars & allies, swifts, and hummingbirds). However, the proportion of names reconciled was reasonably uniform across orders, with outliers in some very small orders where few taxonomic changes have a dramatic effect on proportion (Rheiformes, 2 species; Casuariiformes, 4 species; Suliformes, 10 species), and a few orders which have retained relative taxonomic stability over the past 30 years (e.g. Trogoniformes, Galbuliformes, Ciconiiformes). We also broke down taxonomic reconciliation by family in the large order Passeriformes, where similar patterns emerged. However, in passerines a few large families exhibited high proportions of reconciled names. Speciose passerine families with high proportions of reconciled names included: Phylloscopidae (52%), Leiothrichidae (50%), Sylviidae *sensu stricto* (46%), Scotocercidae (42%), Pellorneidae (41%), Locustellidae (37%), and Timaliidae (27%).

**Fig 2.**
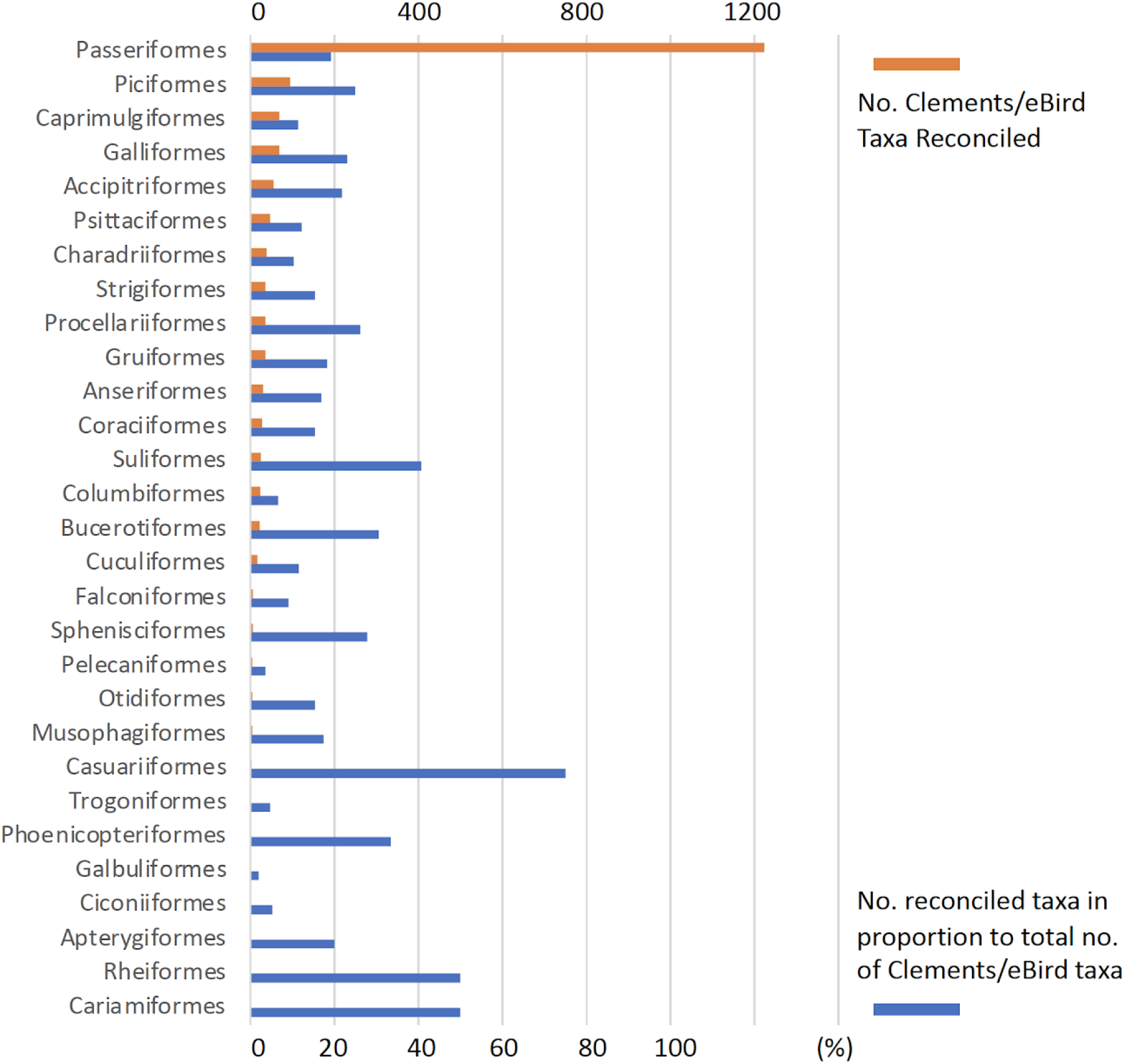
Number and proportion of taxonomic reconciliations applied to GenBank taxa names, by avian order. Note that orders with zero taxonomic reconciliation are not included in the graph.

**Fig 3.**
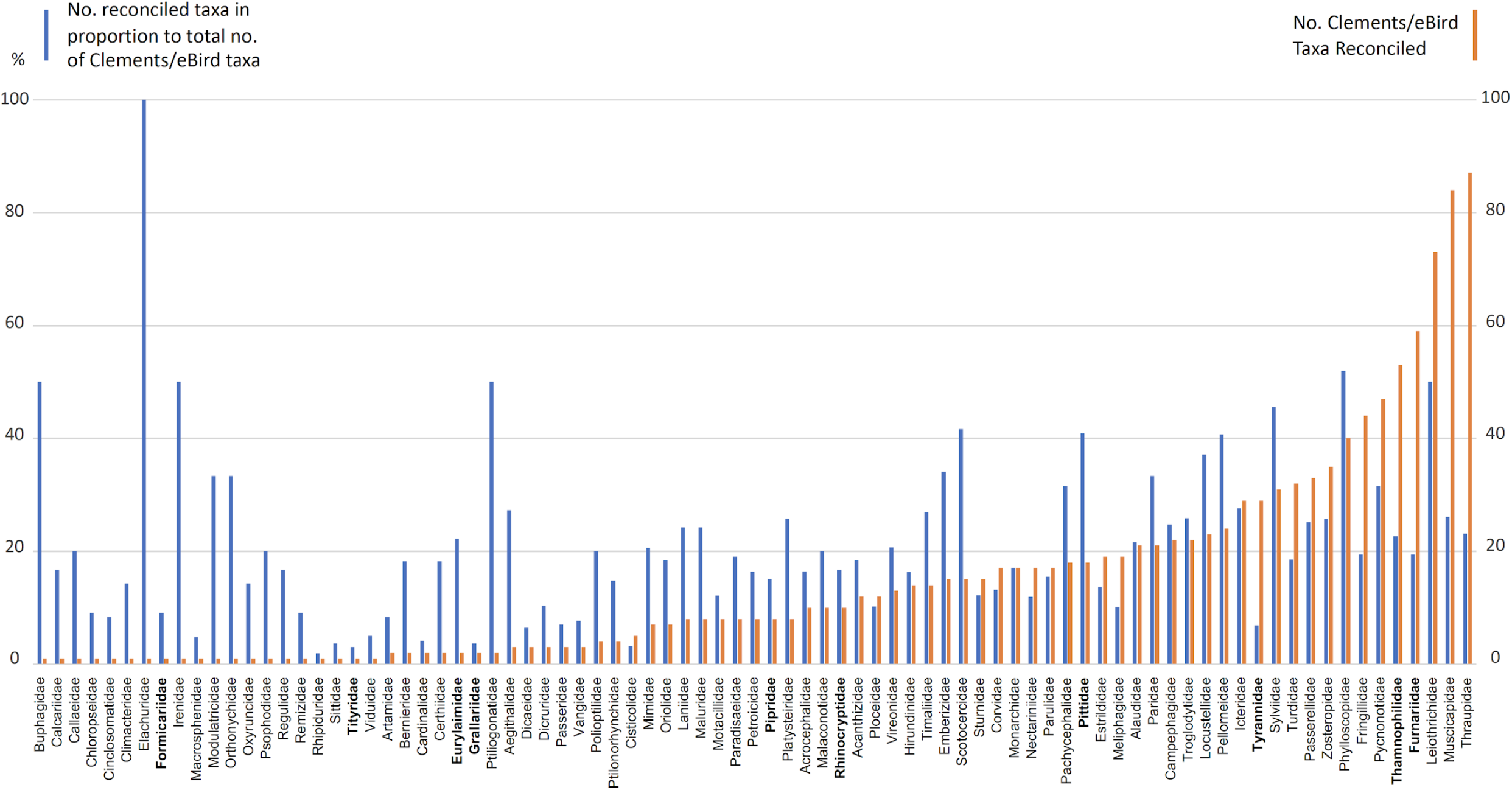
Number and proportion of taxonomic reconciliations applied to GenBank taxa names, by avian family within Passeriformes. Note that families with zero taxonomic reconciliation are not included in the graph.

### Taxonomic reconciliation in relation to IUCN conservation status and geography

There was little relationship between IUCN status and the proportion of taxa reconciled (Table 3; Data Repository D12). The categories Least Concern (LC) Near-threatened (NT) Vulnerable, (V), and Endangered all had similar proportions of taxa reconciled. Critically Endangered taxa were more likely to have had exact matches between GenBank and eBird Clements. Taxa not assessed by IUCN were far less likely to have an exact match between GenBank and eBird/Clements.

**Table 3.** Number and proportion of reconciliations by conservation status and their associated GenBank nucleotide data. The category Extinct includes both extinct taxa and the taxa that were extinct in the wild.

| IUCN conservation status | Num. GenBank nucleotide sequences | Num. eBird/Clements taxa | Num. eBird/Clements taxa reconciled | Proportion eBird/Clements taxa reconciled |
| --- | --- | --- | --- | --- |
| Least Concern | 5277924 | 7976 | 1426 | 17.88% |
| Vulnerable | 782643 | 750 | 113 | 15.07% |
| Near Threatened | 397032 | 932 | 150 | 16.09% |
| Endangered | 259720 | 423 | 70 | 16.55% |
| Critically<br>Endangered | 54789 | 204 | 22 | 10.78% |
| Extinct | 482 | 160 | 4 | 2.50% |
| Data Deficient | 163 | 45 | 7 | 15.56% |
| Not Assessed | 150901 | 286 | 80 | 27.97% |

There was marked geographic variation in the percent of taxa that needed reconciliation. The percentages of widespread taxa (19%) vs. those endemic to one of the realms we considered (18%) were virtually identical (Data Repository D13). Antarctica had no reconciled names, no doubt reflecting the very limited number of taxa found there; the three New World realms and the Australasian realm had the lowest percentages of reconciled names (15% for the North American realm to 17% for the South American realm, Fig. 4; Data Repository D13). Oceanic realms had the highest percentages (up to 37% for the Atlantic ocean; Fig. 4; Data Repository D13).

**Figure 4.**
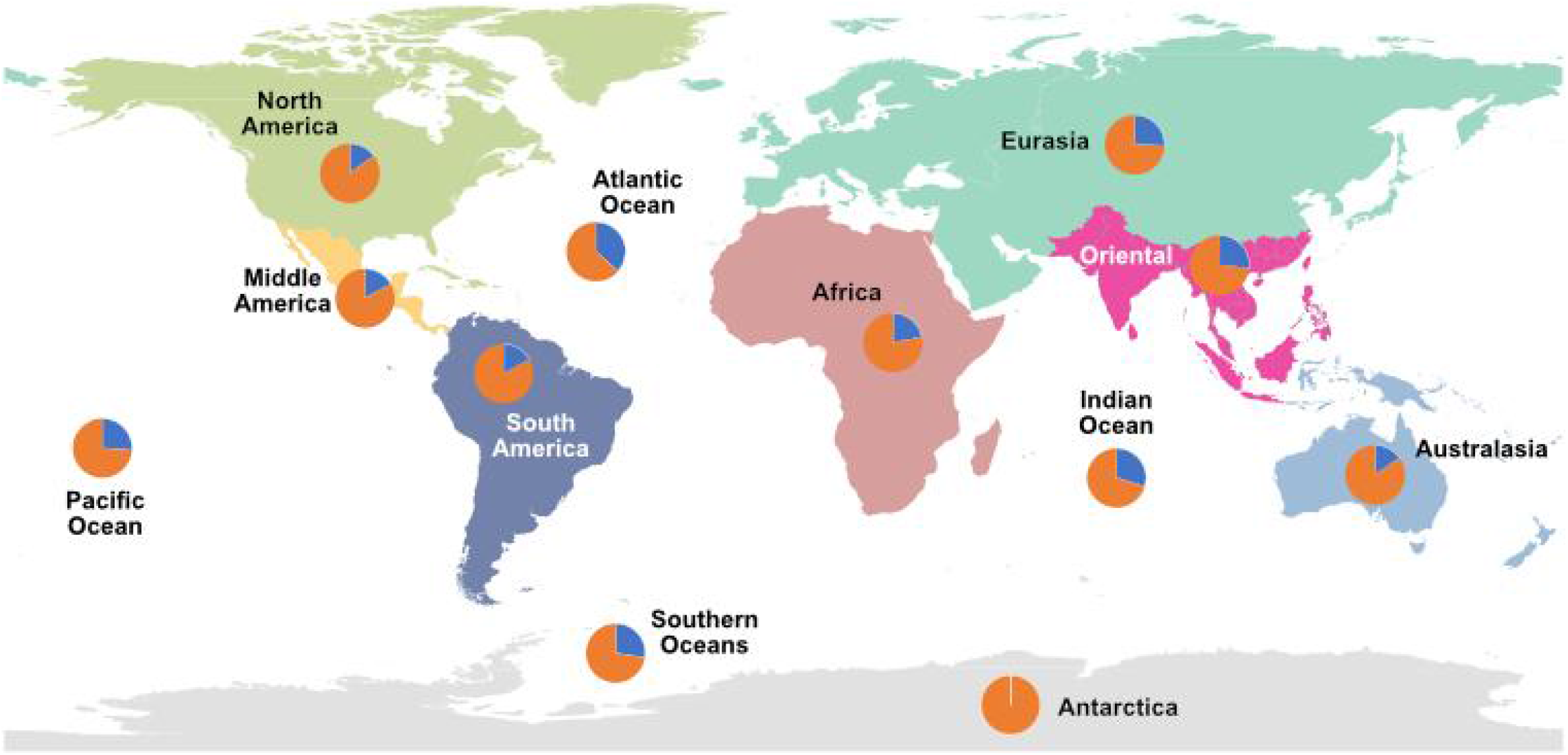
Geographic distribution of taxonomic reconciliations applied to GenBank TaxIDs. With the exception of the Antarctic realm, where there were no reconciliations between GenBank and eBird/Clements, the proportion of reconciled names (blue) ranged from 15% (North America) to 37% (Atlantic Ocean). There is overlap among the individual realms for widespread species.

### Descriptive statistics linking GenBank names to global avian data sources

To assess benefits that reconciling NCBI names with a standardized taxonomy has for the linking of sequence data with phenotypic data, we examined a reconciliation between the Xeno-canto avian sound database (which uses the IOC World Bird list) and the eBird/Clements names. We matched all Xeno-canto avian taxa to eBird/Clements names, except for 13 undescribed and three extinct taxa that are not included in the eBird/Clements 2019 list. 10,166 (93%) of the Xeno-canto names directly matched to eBird/Clements names, and 9506 of those names have available sound recordings (Data Repository D14). The remaining 727 (7%) taxa were reconciled to eBird/Clements using Avibase taxonomic concepts. After reconciliation, we found 9961 eBird/Clements species had sound recordings in Xeno-canto. In Macaulay Library, there are 9609 species with sound data, with an overlap of 9399 species to Xeno-canto. By reconciling GenBank names with eBird/Clements taxonomy, we could easily link sequence data with the two largest avian sound databases which utilize standardized avian taxonomies (Table 4).

**Table 4.** Linking reconciled GenBank names with the two largest avian sound databases which utilize standardized avian taxonomies.

| Database | eBird/Clements<br>species with sound<br>data | Species with<br>both sound and<br>nucleotide data | Sound<br>data<br>only | Nucleotide<br>data only | Neither sound<br>nor nucleotide<br>data |
| --- | --- | --- | --- | --- | --- |
| Xeno-canto | 9961 (92.9%) | 8693 (81.1%) | 1268 | 442 | 318 |
| Macaulay<br>Library | 9609 (89.6%) | 8409 (78.4%) | 1200 | 712 | 386 |
| Combined | 10171 (94.9%) | 8837 (82.4%) | 1333 | 298 | 252 |

### Open source access to taxonomic reconciliation

Our taxonomic reconciliation “RANT: reconciling avian NCBI taxonomy” is open source, and available at GitHub (https://github.com/ebraun68/RANT). Currently, the reconciliation is available for eBird/Clements version 2019 (Data Repository D11). Our intention is to update the reconciliation corresponding to eBird/Clements annual updates.

## Discussion

### Successful linkage of GenBank names to standardized lists

Our reconciliation procedures have successfully linked GenBank taxonomic names (TaxIDs) with avian species and subspecies names regulated by eBird/Clements. Nearly a tenth of all GenBank core nucleotide sequences had a name unrecognized in the eBird/Clemments list, amounting to a total of over 600,000 nucleotide sequences. Hence, it is now easier to link genetic data associated with GenBank TaxIDs to natural populations for comparative work, at least when comparative data have also been reconciled to the eBird/Clements taxonomic lists. If not, these GenBank TaxIDs can still be reconciled to other standardized lists (IOC, BirdLife International, Howard & Moore) through existing resources, namely Avibase and list comparisons freely available from the IOC World Bird List. If avian comparative data do not follow the names of one of these standardized global bird lists, then we strongly advocate that database providers and curators reconcile their aggregated data to one of these standardized lists before its further use and publication.

### Identifying patterns and biases in naming reconciliation

Reconciling GenBank TaxIDs to eBird/Clements names illustrate that naming problems are found throughout the avian tree of life; yet they are concentrated in certain taxonomic groups. Unsurprisingly, these groups tend to have long histories of taxonomic instability. Reconciliation was especially frequent among members of the traditional “Old World Warbler” (Sylviidae *sensu lato*) and “babbler” families (Timaliidae *sensu lato*). These groups have been split into a myriad of smaller families, each of which have undergone substantial revision (Alström et al., 2018, 2011; Cai et al., 2019; Cibois et al., 2002, 1999; Fregin et al., 2012; Moyle et al., 2012).

Outside the Old World warblers and babblers, several other passerine families had high proportions of reconciled names. Forty-one percent of Pittidae names required reconciliation. Perhaps this was because traditionally all pitta species were included in the genus *Pitta*. However, pitta diversity is now divided nearly equally among three genera: *Pitta*, *Hydronis*, and *Erythropitta* (Harvey et al., 2020; Irestedt et al., 2006). Additionally, the highly polytypic *Erythropitta erythrogaster* has been split into 12 species (Irestedt et al., 2013). Another group with a highly polytypic species is the Pachycephalidae, which contains *Pachycephala pectoralis,* which was previously the world’s most polytypic bird species (Andersen et al., 2014; Jonsson et al., 2014). Taxonomic revisions have since split the *P. pectoralis* complex into ~15 species. Reconciliations in Pittidae and Pachycephalidae illustrate how only a few major taxonomic revisions can create stark differences between names used on GenBank and those used in standardized avian bird lists.

One large family, Tyrannidae, had relatively few reconciliations. The eBird/Clements list currently considers 422 species of Tyrannidae, yet the proportion of reconciled names was low, only 7%. Small scale molecular studies have revised parts of tyrannid nomenclature (Hosner and Moyle, 2012; Rheindt et al., 2015). Yet until recently (Harvey et al., 2020; Ohlson et al., 2020), the Tyrannidae has lacked more comprehensive published molecular phylogenies and associated major taxonomic revisions. With the support of these recent publications, we expect the relative nomenclatural stability in Tyrannidae will prove short-lived, and a series of proposed changes will take effect in the coming years.

In addition to taxonomic biases, RANT identified large-scale geographic differences in GenBank name reconciliation. Widespread species— those found in more than one geographic realm, were only slightly more likely to have been subject to taxonomic reconciliation than those limited to a single geographic realm. North America and Australasia proportionally had the fewest reconciled names (Figure 4, Table S1). Both of these realms are comparatively well-studied, so a lack of taxonomic effort is not a viable explanation for their relative stability. One explanation for relative stability in North America and Australia could be the lack of highly problematic groups inhabiting those realms. Very few or no members of taxonomically problematic groups such as Leiothrichidae, Phylloscopidae, or Sylviidae occur in North America or Australia. Although far more diverse than North America or Australia, Middle and South America had only slightly greater proportions of reconciled names (Figure 4), though several of the megadiverse Neotropical families, namely Thraupidae, Furnariidae, and Thamnophilidae were among the families with the greatest total number of reconciliations. Proportionally Old World realms had the most name changes of the terrestrial realms (Figure 4). We suspect the high proportion of reconciled names is related to the concentration of taxonomically problematic groups in these regions, especially the Sylviodea.

The oceanic realms all featured relatively large proportions of reconciled names. Seabirds in particular have undergone extensive taxonomic revisions, driven mostly by molecular genetic work which has revealed great levels of cryptic genetic diversity among ocean basins, breeding islands, and archipelagos (Pyle et al., 2011; Taylor et al., 2019). Among orders, the obligate seafaring groups Procellariiformes, Suliformes, and Sphenisciformes all had large proportions of reconciled names.

Previous authors have raised alarms regarding how taxonomic instability can hamper conservation efforts (Garnett and Christidis, 2017). However, we found that IUCN red listed species were not more likely to have had name reconciliation compared to non-threatened taxa. Among IUCN conservation status categories, “Least Concern” had the greatest proportion of reconciled names whereas “Critically Endangered” had the lowest. Most critically endangered birds are highly range-restricted, and hence are not likely to have been subject to taxonomic splits into multiple species. Taxa not assessed by IUCN had a large proportion of reconciled names, probably driven by the fact that eBird/Cements names not assessed by IUCN are the result of nomenclatural differences between these sources.

### The problem of name application for GenBank sequences

One glaring problem linking taxonomic names to DNA sequences remains, and that is far more insidious than the main problem addressed here. A GenBank TaxID associated with a eBird/Clements name does not necessarily mean that the DNA sequences ascribed to that name will apply correctly (Schoch et al., 2020). Before phylogenetic or population genetic analyses can commence, the correct application of eBird/Clemments names to individual sequences must be verified, a process that is time-consuming and challenging to automate. Below is an example of how the verification process may proceed, drawn from an example of nucleotide data published on GenBank.

The *Robsonius* ground warblers have a complex taxonomic history which highlights many of the naming challenges inherent when working with GenBank data. Originally described in the wren-babbler genus *Napothera* (Rand and Rabor, 1967), for most of its history it has been considered a single species. In 2006 it was split based on new morphological evidence and moved to a new genus (Collar, 2006), and in 2013 a third species was described following the collection of the first adult specimen of true *R. rabori* (Hosner et al., 2013). All four standardized world lists currently recognize all three species: *Robsonius rabori*, *R. sorsogonensis*, and *R. thompsoni*. But Genbank nucleotide data are ascribed to only two TaxIDs: *Robsonius rabori* (TaxID: 1149667, n = 76) and *R. thompsoni* (TaxID: 2162877, n = 3). Most of these data were uploaded under the name *Napothera rabori* prior to use of the *Robsonius* or the epithets *sorsogonensis/thompsoni* in GenBank taxonomy, but sequences are actually derived from all three *Robsonius* species. After tracking down voucher numbers and metadata from publications and voucher specimens, the true taxonomic breakdown of nucleotide sequences is: *Robsonius rabori n =* 7, *R. sorsogonensis n =* 24, and *R. thompsoni n =* 48. Without confirming the application of names, several errors would hamper the use and interpretation of these data. A user might incorrectly conclude that no nucleotide data exist for *R. sorsogonensis*, because its sequences are labeled as *R. rabori*. A user might incorrectly conclude that *R. thompsoni* and *R. rabori* are not genetically distinct, because many *R. thompsoni* sequences are labeled as *R. rabori*. A user might incorrectly conclude that *R. rabori* has exceptional genetic diversity despite its tiny distribution because divergent *R. sorsogonensis* and *R. thompsoni* sequences are each labeled as *R. rabori*.

Resolving name application will be a far more difficult problem to solve than name reconciliation. Name reconciliation requires a set of non-standardized names, a standardized list, and tools or literature to match the non-standardized names to their standardized counterparts. Resolving name application, as in the *Robsonius* example above, requires individual sequence metadata, metadata that is often not recorded in GenBank. Most name application issues arise from splits, when an inclusive former name is applied erroneously to one or more populations with which they were formerly considered conspecific. The most rigorous method to solve these taxonomic problems is to consult voucher specimens to confirm sequence identity. However, many GenBank sequences lack proper voucher specimen information (Buckner et al., 2021; Peterson et al., 2007), as we also noted. After filtering the “Aves” sequences in the nucleotide database to include only genomic DNA/RNA nucleotide sequences (excluding mRNA or rRNA sequences) from the INSDC (GenBank, not RefSeq) source database, we estimated only 17% (484,232) of the 2,902,805 sequences included voucher information anywhere in the full GenBank record. While some other samples may have information included that could be used to trace the source of the sample, it is clear that the majority of available sequence data lacks such information. While there is now a GenBank voucher field, it is not required, and is easy for sequence authors to omit. Sometimes voucher information can also be found in the sequence definition line. While checking vouchers one-by-one is not feasible for large scale metadata correction of what, at present count, is over six million avian nucleotide sequences, it can still be used to resolve at least some problems.

Aside from vouchers, locality metadata is useful when resolving name application problems. Latitude and longitude can be included in GenBank metadata, but often it is not. In some cases, it can be found in published papers or their online supplements, or in publicly shared museum databases if the samples were properly vouchered and digitized. However, like checking specimen vouchers, this laborious task is not suited to large-scale applications. However, automated georeferencing algorithms may be a viable tool to improve sequence attribution geographically (Miraldo et al., 2016). When splits apply to allopatric populations, the latitude/longitude of the sample origin solves name application. However, when these splits do not apply cleanly to allopatric populations, or when migratory populations of split taxa overlap for part of the year, further information will be needed to resolve sequence identity.

### A call for expert curation of avian GenBank sequence metadata

RANT is a first step towards active and decentralized management of metadata associated with avian sequence data. These standardized names provide a new benchmark for managing large-scale sequence meta-analyses, but many data problems remain--- particularly the challenge of verifying names application to individual DNA sequences. Although GenBank provides a vast and important resource, large biodiversity datasets need constant management and expert curation to maximize their usefulness (Sangster and Luksenburg, 2021; Schoch et al., 2020). One solution is to maintain a parallel database to update and store metadata related to GenBank sequences, but free of its restrictive updating policies (Riginos et al., 2020). With such a system, a team of expert curators could gather, review, proofread, and provide supplemental metadata associated with GenBank sequences (Marques et al., 2013), linked to the actual sequence data housed at GenBank through the accession number. In addition to validating metadata, curators can permanently flag or provide feedback on potential problematic sequences (De Silva et al., 2019; Sangster and Luksenburg, 2021). These strategies are effective for curating far larger sets of biodiversity data collected largely by non-professional scientists (Robertson et al., 2014; Sullivan et al., 2009).

## ACKNOWLEDGEMENTS

We will thank anonymous reviewers who provided comments that improved the manuscript.

---

**Supplementary Table S1.**
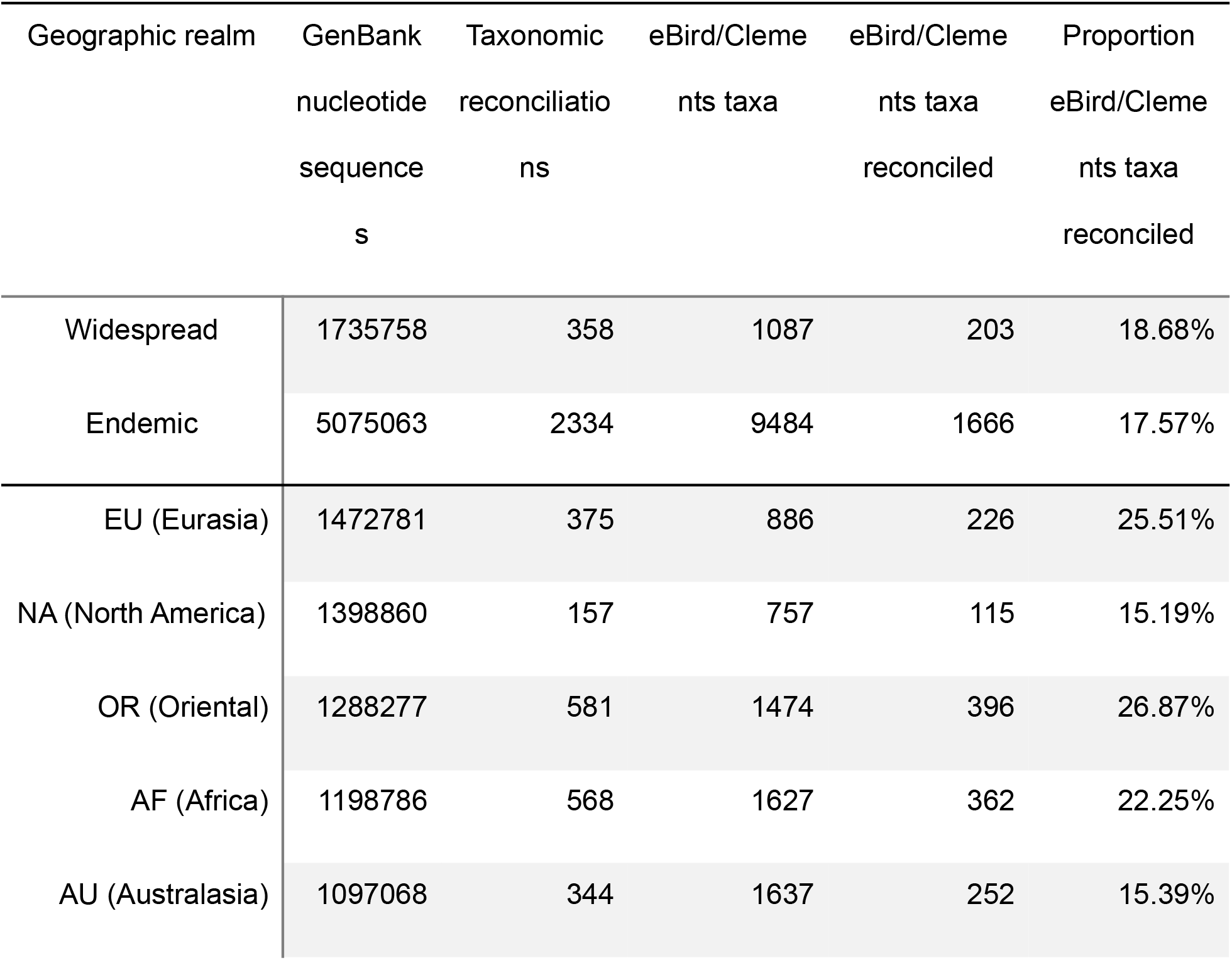
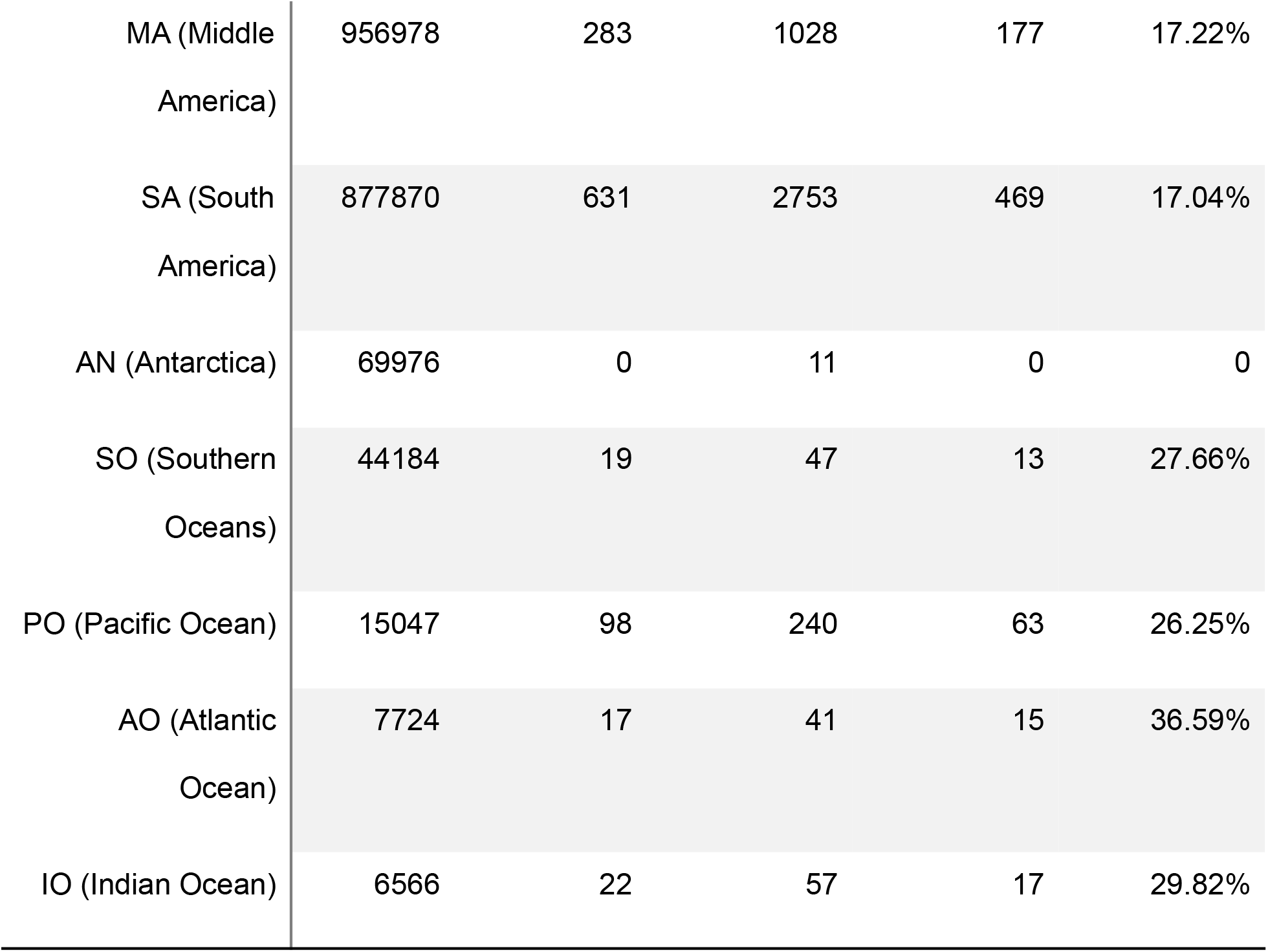
Number and proportion of reconciliations by biogeographical realm and their associated GenBank nucleotide data. “Widespread” indicates taxa that occur in two or more realms listed below, and “Endemic” indicates taxa that occur in only one realm. There is overlap among the individual realms for widespread taxa.

